# Small extracellular vesicle-enriched preparations derived from genetically modified multipotent mesenchymal stromal cells promote fibroblast proliferation and extracellular matrix remodeling in vitro

**DOI:** 10.64898/2026.09.17.752298

**Authors:** Oleg G. Makeev, Artem V. Korotkov, Angelina A. Seliverstova

## Abstract

Impaired skin repair is associated with insufficient fibroblast activity and dysregulated extracellular matrix turnover. Extracellular vesicle-enriched preparations derived from multipotent mesenchymal stromal cells (MMSCs) may provide a cell-free approach to modulate these processes; however, the effects of preparations from genetically modified producer cells remain incompletely characterized. We evaluated preparations obtained from native MMSCs and from MMSCs separately transfected with OCT4/SOX2, HIF-1alpha, or Klotho expression plasmids. A 1:1:1 mixture of preparations from the three modified cell populations was examined as a combined treatment. Rat skin fibroblasts were exposed to the preparations for 24, 48, or 72 h. Fibroblast density and concentrations of matrix metalloproteinase-1 (MMP-1), matrix metalloproteinase-3 (MMP-3), collagen type I, and collagen type III were assessed. The combined preparation produced the largest observed increase in fibroblast density, reaching 112.63 +/−5.55 × 10^3 cells/cm2 at 72 h, compared with 40.60 +/−1.12 × 10^3 cells/cm2 for the native-MMSC preparation and 37.10 +/−3.01 × 10^3 cells/cm2 in the untreated control. At 24 h, MMP-1 and MMP-3 concentrations in the combined group were 140.0 +/−4.77 ng/mL and 144.0 +/−4.74 ng/mL, respectively, compared with 283.1 +/−25.2 ng/mL and 208.1 +/−39.2 ng/mL in controls. Collagen type III concentration increased to 1.95 +/−0.25 ng/mL at 72 h versus 1.15 +/−0.15 ng/mL in controls. These findings indicate that the combined preparation was associated with enhanced fibroblast proliferation and altered extracellular matrix remodeling in vitro. Interpretation is limited by triplicate technical replication, volume-based dosing, and incomplete vesicle characterization.

## Introduction

Impaired repair of skin injuries is associated with persistent inflammation, insufficient restoration of the fibroblast population, and dysregulated extracellular matrix turnover. Chronic wounds, burns, and soft-tissue defects may progress to delayed closure and formation of functionally inferior scar tissue. Improving the regulation of fibroblast growth and matrix remodeling remains an important objective in regenerative medicine.

Extracellular vesicles released by multipotent mesenchymal stromal cells have attracted attention as potential cell-free mediators of intercellular communication. Their molecular contents can influence recipient-cell proliferation, inflammatory signaling, and extracellular matrix homeostasis. Genetic modification of producer cells may alter the biological properties of their secreted extracellular-vesicle-enriched preparations and could provide a route to more targeted regenerative interventions.

OCT4 and SOX2 are associated with cellular plasticity and partial reprogramming programs. HIF-1alpha is a central mediator of cellular adaptation to hypoxia and has been linked to tissue-repair responses. Klotho has been implicated in pathways relevant to oxidative stress, fibrosis, and cellular aging. The combined influence of extracellular-vesicle-enriched preparations obtained from MMSCs modified with these constructs on fibroblast proliferation and matrix-associated outcomes remains insufficiently defined.

The aim of this study was to evaluate the effects of extracellular-vesicle-enriched preparations derived from native and genetically modified MMSCs on fibroblast proliferation and extracellular matrix remodeling in vitro.

## Materials and methods

### Study design

This was a controlled, non-randomized in vitro experiment with six parallel treatment groups. Outcomes were assessed at 24, 48, and 72 h. Each group at each time point was assessed in three technical wells. Results are presented as mean +/−standard error of the mean (SEM) across technical replicates.

### Animals and ethical approval

Ten adult male Wistar rats, approximately 20 weeks old and weighing approximately 180 g, were used as tissue donors. All animal procedures were conducted in accordance with internationally accepted principles for the humane treatment of laboratory animals and institutional requirements. The study was approved by the Local Ethics Committee of Ural State Medical University, Yekaterinburg, Russian Federation (Protocol No. 5, 24 June 2015). No human participants or human-derived materials were involved.

### Isolation and culture of MMSCs

MMSCs were isolated by enzymatic processing of the stromal vascular fraction from abdominal-wall adipose tissue of Wistar rats. Cells were expanded in DMEM/F12 supplemented with 10% fetal bovine serum under standard culture conditions. Second-passage MMSCs were used for the study. MMSC identity was assessed by CD-marker expression, including CD73, CD90, and CD105, with minimal expression of CD34, CD45, CD14, and CD19.

### Fibroblast culture

Skin fibroblasts were obtained from rat tail skin after euthanasia under thiopental anesthesia by decapitation. Cells were cultured at 37 C in a humidified atmosphere containing 5% CO2 and seeded at 1 × 10^4 cells/cm2.

### Genetic modification of MMSC producer cells

MMSCs were separately transfected with OCT4/SOX2 (Addgene plasmid #6945), HIF-1alpha (Addgene plasmid #105108), or Klotho (Addgene plasmid #17712) expression constructs using Escort III (Sigma-Aldrich) liposomal complexation. Transfection efficiency was assessed by RT-PCR. Detailed RT-PCR conditions and quantitative transfection results were not available for this report.

### Isolation and characterization of extracellular-vesicle-enriched preparations

Conditioned medium from native and genetically modified MMSC cultures was processed using a combined workflow that included centrifugation at 600 g for 60 min, centrifugation at 15,000-17,000 g for 220 min, ultracentrifugation at 100,000 g for 300 min, filtration through a 0.22-um filter, polyethylene glycol precipitation at 4 C for 24 h, and a subsequent ultracentrifugation step at 100,000 g for 120 min. The final pellet was resuspended in 450 uL phosphate-buffered saline and stored at −20 C until use.

Transmission electron microscopy and immunoaffinity ELISA capture were used for preparation identification. Quantitative particle-size distribution, particle concentration, molecular-marker panel results, rotor details, temperature during centrifugation steps, and starting conditioned-medium volumes were not available. Accordingly, the preparations are described as extracellular-vesicle-enriched preparations rather than as purified exosomes.

### Experimental groups and cell treatment

Rat skin fibroblasts were assigned to six groups: (1) untreated control, which received no added preparation; (2) preparation derived from native MMSCs; (3) preparation from OCT4/SOX2-transfected MMSCs; (4) preparation from HIF-1alpha-transfected MMSCs; (5) preparation from Klotho-transfected MMSCs; and (6) a combined preparation consisting of the three genetically modified preparations mixed in a 1:1:1 ratio.

Fibroblast cultures in groups 2-6 received 20 uL per well of the respective extracellular-vesicle-enriched preparation. The untreated control received no additional volume. No particle- or protein-based normalization of the preparations was documented.

### Outcome measures

Fibroblast density was measured at 24, 48, and 72 h. Concentrations of MMP-1, MMP-3, collagen type I, and collagen type III were measured in culture media using commercial ELISA assays from Cloud-Clone Corp. Product catalog numbers, assay ranges, and documented species-validation details were not available.

### Statistical analysis

Statistical analysis was conducted in RStudio version 1.1.419. Distribution normality was assessed using the Shapiro-Wilk test. Student t tests and Mann-Whitney U tests were used as appropriate, with p < 0.05 considered statistically significant. The data available for this report did not include raw values, exact p values, or information on multiple-comparison adjustment. Because the reported triplicates were technical replicates, statistical inference should be interpreted cautiously and does not demonstrate independent biological reproducibility.

## Results

### Fibroblast proliferation

All tested preparations increased fibroblast density to varying extents. The largest observed response occurred in the combined-treatment group. At 24 h, fibroblast density was 24.53 +/−2.21 × 10^3 cells/cm2 in the combined group, compared with 5.71 +/−0.96 × 10^3 cells/cm2 in the untreated control and 11.00 +/−1.12 × 10^3 cells/cm2 in the native-MMSC preparation group. At 48 h, the respective values were 62.57 +/−3.97, 25.63 +/−1.36, and 30.25 +/−1.34 × 10^3 cells/cm2. At 72 h, fibroblast density reached 112.63 +/−5.55 × 10^3 cells/cm2 in the combined group, compared with 37.10 +/−3.01 × 10^3 cells/cm2 in the untreated control and 40.60 +/−1.12 × 10^3 cells/cm2 in the native-MMSC preparation group.

At 72 h, the HIF-1alpha group reached 81.23 +/−6.39 × 10^3 cells/cm2 and the Klotho group reached 63.22 +/−4.87 × 10^3 cells/cm2. The OCT4/SOX2 group reached 47.78 +/−3.01 × 10^3 cells/cm2.

### MMP-1 and MMP-3 concentrations

MMP-1 concentration was lower at early time points in the OCT4/SOX2 and combined groups. At 24 h, MMP-1 was 150.0 +/−5.50 ng/mL in the OCT4/SOX2 group and 140.0 +/−4.77 ng/mL in the combined group, compared with 283.1 +/−25.2 ng/mL in the untreated control. At 48 h, MMP-1 was 166.0 +/−5.68 ng/mL in the OCT4/SOX2 group and 153.0 +/−6.68 ng/mL in the combined group, compared with 233.7 +/−20.3 ng/mL in the untreated control. By 72 h, differences between groups were smaller.

MMP-3 showed a similar early pattern. At 24 h, MMP-3 concentration was 160.0 +/−4.50 ng/mL in the OCT4/SOX2 group and 144.0 +/−4.74 ng/mL in the combined group, compared with 208.1 +/−39.2 ng/mL in the untreated control. At 48 h, MMP-3 was 161.0 +/−5.60 ng/mL in the OCT4/SOX2 group and 158.0 +/−7.68 ng/mL in the combined group, compared with 211.1 +/−30.3 ng/mL in the untreated control. At 72 h, differences were not reported as statistically significant.

### Collagen type I and type III concentrations

Collagen type I values changed modestly. In the combined group, concentrations were 17.99 +/−0.12 ng/mL at 24 h, 18.53 +/−0.27 ng/mL at 48 h, and 19.41 +/−0.10 ng/mL at 72 h. The 72-h value was reported as significantly different from the control value of 17.95 +/−0.15 ng/mL.

Collagen type III showed a more pronounced response in the combined group. At 48 h, its concentration was 1.23 +/−0.13 ng/mL compared with 0.79 +/−0.10 ng/mL in the untreated control. At 72 h, collagen type III reached 1.95 +/−0.25 ng/mL in the combined group, compared with 1.15 +/−0.15 ng/mL in the untreated control.

**Table 1.** Fibroblast density (× 10^3 cells/cm2, mean +/−SEM; n = 3 technical wells)

| Time, h | Untreated control | Native MMSC preparation | OCT4/SOX2 preparation | HIF-1alpha preparation | Klotho preparation | Combined preparation |
| --- | --- | --- | --- | --- | --- | --- |
| 24 | $5.71 \pm 0.96$ | $11.00 \pm 1.12^*$ | $22.07 \pm 4.10^{**}$ | $19.98 \pm 1.97^*$ | $7.88 \pm 0.98$ | $24.53 \pm 2.21^{**}$ |
| 48 | $25.63 \pm 1.36$ | $30.25 \pm 1.34^*$ | $35.11 \pm 3.38^{**}$ | $34.45 \pm 1.54^*$ | $35.42 \pm 2.99^{**}$ | $62.57 \pm 3.97^{**}$ |
| 72 | $37.10 \pm 3.01$ | $40.60 \pm 1.12$ | $47.78 \pm 3.01^{**}$ | $81.23 \pm 6.39^{**}$ | $63.22 \pm 4.87^{**}$ | $112.63 \pm 5.55^{**}$ |

\* p < 0.05 versus untreated control. \*\* p < 0.05 versus native MMSC preparation, as reported in the source dataset.

**Table 2.** MMP-1 concentration in culture medium (ng/mL, mean +/−SEM; n = 3 technical wells)

| Time, h | Untreated control | Native MMSC preparation | OCT4/SOX2 preparation | HIF-1alpha preparation | Klotho preparation | Combined preparation |
| --- | --- | --- | --- | --- | --- | --- |
| 24 | $283.1 \pm 25.2$ | $203.1 \pm 22.2$ | $150.0 \pm 5.50^*$ | $200.3 \pm 19.2$ | $198.9 \pm 19.9$ | $140.0 \pm 4.77^*$ |
| 48 | $233.7 \pm 20.3$ | $227.1 \pm 21.3$ | $166.0 \pm 5.68^*$ | $202.1 \pm 20.3$ | $200.1 \pm 20.6$ | $153.0 \pm 6.68^*$ |
| 72 | $204.2 \pm 23.6$ | $201.1 \pm 22.0$ | $197.8 \pm 6.01$ | $200.1 \pm 18.0$ | $200.5 \pm 19.1$ | $192.8 \pm 6.91$ |

\* p < 0.05 versus untreated control, as reported in the source dataset.

**Table 3.** MMP-3 concentration in culture medium (ng/mL, mean +/−SEM; n = 3 technical wells)

| Time, h | Untreated control | Native MMSC preparation | OCT4/SOX2 preparation | HIF-1alpha preparation | Klotho preparation | Combined preparation |
| --- | --- | --- | --- | --- | --- | --- |
| 24 | $208.1 \pm 39.2$ | $199.2 \pm 21.2$ | $160.0 \pm 4.50^*$ | $201.8 \pm 18.0$ | $199.9 \pm 29.9$ | $144.0 \pm 4.74^*$ |
| 48 | $211.1 \pm 30.3$ | $200.8 \pm 31.7$ | $161.0 \pm 5.60^*$ | $202.9 \pm 19.3$ | $200.0 \pm 30.6$ | $158.0 \pm 7.68^*$ |
| 72 | 209.1 +/- 33.6 | 205.1 +/- 32.8 | 187.3 +/- 6.11 | 210.1 +/- 17.6 | 202.7 +/- 19.0 | 188.1 +/- 7.91 |

\* p < 0.05 versus untreated control, as reported in the source dataset.

**Table 4.** Collagen type I concentration in culture medium (ng/mL, mean +/−SEM; n = 3 technical wells)

| Time, h | Untreated control | Native MMSC preparation | OCT4/SOX2 preparation | HIF-1alpha preparation | Klotho preparation | Combined preparation |
| --- | --- | --- | --- | --- | --- | --- |
| 24 | 16.75 +/- 0.11 | 16.69 +/- 0.12 | 16.88 +/- 0.18 | 16.78 +/- 0.18 | 16.98 +/- 0.14 | 17.99 +/- 0.12 |
| 48 | 16.78 +/- 0.10 | 16.87 +/- 0.13 | 16.78 +/- 0.12 | 16.68 +/- 0.17 | 17.33 +/- 0.20 | 18.53 +/- 0.27 |
| 72 | 17.95 +/- 0.15 | 17.25 +/- 0.13 | 17.97 +/- 0.16 | 17.25 +/- 0.15 | 17.21 +/- 0.12 | 19.41 +/- 0.10* |

\* p < 0.05 versus untreated control, as reported in the source dataset.

**Table 5.** Collagen type III concentration in culture medium (ng/mL, mean +/−SEM; n = 3 technical wells)

| Time, h | Untreated control | Native MMSC preparation | OCT4/SOX2 preparation | HIF-1alpha preparation | Klotho preparation | Combined preparation |
| --- | --- | --- | --- | --- | --- | --- |
| 24 | 0.75 +/- 0.01 | 0.77 +/- 0.01 | 0.73 +/- 0.01 | 0.75 +/- 0.01 | 0.70 +/- 0.01 | 0.76 +/- 0.01 |
| 48 | 0.79 +/- 0.10 | 0.76 +/- 0.10 | 0.78 +/- 0.10 | 0.74 +/- 0.09 | 0.74 +/- 0.09 | 1.23 +/- 0.13* |
| 72 | 1.15 +/- 0.15 | 1.05 +/- 0.15 | 0.95 +/- 0.11 | 1.05 +/- 0.16 | 1.11 +/- 0.09 | 1.95 +/- 0.25* |

\* p < 0.05 versus untreated control, as reported in the source dataset.

## Discussion

In this in vitro study, extracellular-vesicle-enriched preparations derived from genetically modified MMSCs were associated with changes in fibroblast density and matrix-associated outcomes. The combined preparation, formed by mixing preparations from OCT4/SOX2-, HIF-1alpha-, and Klotho-transfected producer-cell populations in equal proportions, produced the largest observed increase in fibroblast density. It was also associated with lower MMP-1 and MMP-3 concentrations at the early time points and with higher collagen type III concentration at later time points.

The results are compatible with the hypothesis that molecular programs associated with partial reprogramming, hypoxia response, and Klotho-related regulation may influence the paracrine activity of MMSC-derived extracellular-vesicle-enriched preparations. However, this study does not establish the molecular contents responsible for the observed outcomes. Nor does the design demonstrate pharmacologic or biologic synergy; accordingly, the findings are described as the largest observed effects of the combined preparation rather than as proof of synergistic action.

The reduced MMP-1 and MMP-3 concentrations observed at 24 and 48 h may indicate altered proteolytic conditions in the culture environment. The increase in collagen type III in the combined group is consistent with an early matrix-remodeling response. These in vitro observations should not be interpreted as evidence of clinical wound-healing efficacy.

Important limitations include the in vitro design, the use of technical rather than independent biological replicates, absence of a volume-matched vehicle in the untreated control, volume-based rather than particle- or protein-based dosing, and incomplete characterization of the extracellular-vesicle-enriched preparations. The mixed centrifugation and polyethylene glycol workflow may co-isolate non-vesicular extracellular components. Consequently, the reported findings should be regarded as preliminary and require independent biological replication, standardized dose normalization, full EV characterization, and validation in relevant in vivo models.

## Conclusion

Extracellular-vesicle-enriched preparations derived from genetically modified MMSCs were associated with increased fibroblast proliferation and changes in extracellular matrix remodeling markers in vitro. The 1:1:1 combined preparation from OCT4/SOX2-, HIF-1alpha-, and Klotho-modified MMSC producer cells produced the largest observed effects under the tested conditions. Given the technical replication, incomplete characterization, and volume-based dosing, these preliminary findings require confirmation in independently replicated and fully characterized experiments.

## Related manuscript statement

This report uses an experimental platform shared with a companion manuscript focused on TERT protein levels in cultured rat skin fibroblasts under the same treatment conditions. The present manuscript reports a distinct outcome set limited to fibroblast density, MMP-1, MMP-3, collagen type I, and collagen type III. No TERT numerical data, tables, or figures are included here. The companion manuscript will disclose this relationship and will not duplicate the outcome data reported in the present manuscript.

## Data availability

The summarized data underlying the results are included in this manuscript. Additional underlying records are available from the corresponding author upon reasonable request, subject to institutional requirements.

## Author contributions

O.G.M. conceived and designed the study, provided scientific supervision, and revised the manuscript. A.V.K. conducted experiments and contributed to data analysis, interpretation, and manuscript preparation. A.A.S. contributed to material collection and processing, laboratory work, literature review, and preparation of the initial manuscript draft. All authors approved the final manuscript and accept responsibility for all aspects of the work.

## Use of generative AI

Generative artificial intelligence was not used to generate primary research results, analyze experimental data, or determine the scientific conclusions of this study. It may have been used for technical language editing; all scientific content was reviewed and approved by the authors.

## References

1. Li D, Chen M, Li W, Xu X, Li Q. Global burden of viral skin diseases from 1990 to 2021: a systematic analysis for the global burden of disease study 2021. Front Public Health. 2025;13:1464372. doi:10.3389/fpubh.2025.1464372.

2. Vecin NM, Kirsner RS. Skin substitutes as treatment for chronic wounds: current and future directions. Front Med (Lausanne). 2023;10:1154567. doi:10.3389/fmed.2023.1154567.

3. Przekora A. A concise review on tissue engineered artificial skin grafts for chronic wound treatment: can we reconstruct functional skin tissue in vitro? Cells. 2020;9(7):1622. doi:10.3390/cells9071622.

4. Riza SM, Porosnicu AL, Cepi PA, Parasca SV, Sinescu RD. Integrating regenerative medicine in chronic wound management: a single-center experience. Biomedicines. 2025;13(8):1827. doi:10.3390/biomedicines13081827.

5. Xie S, Zhang Q, Jiang L. Current knowledge on exosome biogenesis, cargo-sorting mechanism and therapeutic implications. Membranes (Basel). 2022;12(5):498. doi:10.3390/membranes12050498.

6. Welsh JA, Van Der Pol E, Arkesteijn GJA, et al. Minimal information for studies of extracellular vesicles (MISEV2023): From basic to advanced approaches. J Extracell Vesicles. 2024;13(2):e12404. doi:10.1002/jev2.12404.

7. Sadeghi S, et al. Exosomes derived from mesenchymal stem cells: a promising cell-free therapeutic tool for cutaneous wound healing. Biochimie. 2023;209:73–84. doi:10.1016/j.biochi.2023.01.013.

8. Yuan M, et al. Biogenesis, composition and potential therapeutic applications of mesenchymal stem cells derived exosomes in various diseases. Int J Nanomedicine. 2023;18:3177–3210. doi:10.2147/IJN.S407029.

9. Chen S, et al. Bioengineered MSC-derived exosomes in skin wound repair and regeneration. Front Cell Dev Biol. 2023;11:1029671. doi:10.3389/fcell.2023.1029671.

10. Kim KP, Han DW, Kim J, Scholer HR. Biological importance of OCT transcription factors in reprogramming and development. Exp Mol Med. 2021;53(6):1018–1028. doi:10.1038/s12276-021-00637-4.

11. Browder KC, Reddy P, Yamamoto M, et al. In vivo partial reprogramming alters age-associated molecular changes during physiological aging in mice. Nat Aging. 2022;2(3):243–253. doi:10.1038/s43587-022-00183-2.

12. Basheeruddin M, Qausain S. Hypoxia-Inducible Factor 1-Alpha (HIF-1alpha): an essential regulator in cellular metabolic control. Cureus. 2024;16(7):e63852. doi:10.7759/cureus.63852.

13. Ren T, Wen ZK, Liu ZM, et al. A small molecule HIF-1alpha stabilizer that accelerates diabetic wound healing. Nat Commun. 2021;12(1):3363. doi:10.1038/s41467-021-23448-7.

14. Torres-Velarde JM, Allen KN, Su Y, et al. Pathobiology of the Klotho antiaging protein and therapeutic considerations. Front Aging. 2022;3:931331. doi:10.3389/fragi.2022.931331.

15. Jiang C, Fang X, et al. New insights into the role of Klotho in inflammation and fibrosis: molecular and cellular mechanisms. Front Immunol. 2024;15:1454142. doi:10.3389/fimmu.2024.1454142.

